# Air and surface contamination in non-health care settings among 641 environmental specimens of 39 COVID-19 cases

**DOI:** 10.1101/2020.07.09.195008

**Authors:** Lei Luo, Dan Liu, Hao Zhang, Zhihao Li, Ruonan Zhen, Xiru Zhang, Huaping Xie, Weiqi Song, Jie Liu, Qingmei Huang, Jingwen Liu, Xingfen Yang, Zongqiu Chen, Chen Mao

**Affiliations:** Guangzhou Center for Disease Control and Prevention, Guangzhou, China; Department of Epidemiology, School of Public Health, Southern Medical University, Guangzhou, China; Food Safety and Health Research Center, School of Public Health, Southern Medical University, Guangzhou, China

**Keywords:** SARS-CoV-2, COVID-19, Environmental Contamination

## Abstract

**Background:** Little is known about the SARS-CoV-2 contamination of environmental surfaces and air in non-health care settings among COVID-19 cases.

**Methods and findings:** We explored the SARS-CoV-2 contamination of environmental surfaces and air by collecting air and swabbing environmental surfaces among 39 COVID-19 cases in Guangzhou, China. The specimens were tested by RT-PCR testing. The information collected for COVID-19 cases included basic demographic, clinical severity, onset of symptoms, radiological testing, laboratory testing and hospital admission. A total of 641 environmental surfaces and air specimens were collected among 39 COVID-19 cases before disinfection. Among them, 20 specimens (20/641, 3.1%) were tested positive from 9 COVID-19 cases (9/39, 23.1%), with 5 (5/101, 5.0%) positive specimens from 3 asymptomatic cases, 5 (5/220, 2.3%) from 3 mild cases, and 10 (10/374, 2.7%) from 3 moderate cases. All positive specimens were collected within 3 days after diagnosis, and 10 (10/42, 23.8%) were found in toilet (5 on toilet bowl, 4 on sink/faucet/shower, 1 on floor drain), 4 (4/21, 19.0%) in anteroom (2 on water dispenser/cup/bottle, 1 on chair/table, 1 on TV remote), 1 (1/8, 12.5%) in kitchen (1 on dining-table), 1 (1/18, 5.6%) in bedroom (1 on bed/sheet pillow/bedside table), 1 (1/5, 20.0%) in car (1 on steering wheel/seat/handlebar) and 3 (3/20, 21.4%) on door knobs. Air specimens in room (0/10, 0.0%) and car (0/1, 0.0%) were all negative.

**Conclusions:** SARS-CoV-2 was found on environmental surfaces especially in toilet, and could survive for several days. We provided evidence of potential for SARS-CoV-2 transmission through contamination of environmental surfaces.

## Introduction

The Coronavirus Disease 2019 (COVID-19) pandemic has precipitated a global crisis, and it has resulted in 5404512 confirmed cases including with 343514 deaths globally as of May 26, 2020[1]. Reported transmission modes of Severe Acute Respiratory Syndrome Coronavirus 2 (SARS-CoV-2) among humans mainly through respiratory droplets produced when an infected case sneezes or coughs[2]. People may be infected by inhalation of virus laden liquid droplets, and infection is more likely when someone are in close contact with COVID-19 cases[2–4]. However, the importance of indirect contact transmission, such as environmental contamination, is uncertain[5–7]. Evidences suggested that environmental contamination with SARS-CoV-2 is likely to be high, and it is supported by recent researches focus on environmental contamination from COVID-19 cases in hospital[5–8]. Hospitals already have perfect disinfection measures and are less likely to appear super-spreaders compared with community and family[4,9–11]. However, the role of air and surface contamination in non-health care settings is still need to be explored. Therefore, it is vital to understand the environmental contamination of infected cases by SARS-CoV-2 in non-health care settings, which was a vital aspect of controlling the spread of the epidemic.

To address this question, in this study, we sampled total of 641 surfaces environmental and air specimens among 39 cases in Guangzhou, China, to explore the surrounding environmental surfaces and air contamination by SARS-CoV-2 in non-health care settings.

## Methods

### Study design and setting

Based on COVID-19 case reports from Jan 27 to Apr 9, 2020, environmental surfaces and air specimens were collected by Guangzhou CDC (GZCDC) from Feb 6 to Apr 10, 2020. The environmental surfaces specimens of COVID-19 cases sampled in home, hotel, public area, restaurant, marketplace, car and pet, which was associated with COVID-19 cases’ life trajectory before hospitalization. Air specimens of COVID-19 cases also sampled in their room (home or hotel). Based on cases’ reported activity tracks, the number of specimens collected per COVID-19 case varied. All specimens were collected before disinfection.

### Demographic and epidemiological data

Case investigations were collected for COVID-19 cases including basic demographic (age, sex, imported cases or not), date of symptoms onset, date of diagnosis, date of environmental specimens collection, clinical severity, onset of symptoms (fever, dry cough, expectoration, fatigue, myalgia and diarrhea), radiological testing (CT double lung abnormalities), laboratory testing (white blood cell count, lymphocyte, lymphocyte percentage, neutrophilic granulocyte, and neutrophilic granulocyte percentage) and hospital admission.

### Sample Collection

The environmental specimens of SARS-CoV-2 were collected by qualified technicians who had received biosafety training (who had passed the training) and were equipped with corresponding laboratory skills. Personal protective equipment (PPE) were required for sampling personnel including N95 masks or masks with higher filtration efficiency, goggles, protective clothing, double-layer latex gloves and waterproof boot covers[12]. Sterile premoistened swabs were used for sampling, and then were put into a tube containing 2 to 3 mL virus preservation solution (or isotonic saline solution, tissue culture solution, or phosphate buffer) with swabs’ tail discarded and cap tightened. For details, please refer to Protocol for Prevention and Control of COVID-19 (Edition 6) issued by the National Health Commission of the People’s Republic of China[13]. The following types of surfaces were swabbed: (1) toilet (sink, faucet, shower, toilet bowl, floor drain); (2) anteroom (chair, table, TV remote, water dispenser, cup, bottle, TV bench, handrail); (3) kitchen (cutting board, bowl, chopsticks); (4) bedroom (bed, sheet pillow, bedside table, telephone, computer, mouse, air conditioner, fan); (5) door knobs and switch buttons in room; (6) air in room; (7) outside room (elevator, elevator button, stair armrest); (8) car (steering wheel, seat, handlebar, air); (9) pet (mouth, nose, anus). For each case, not all of the above samples were collected, but only samples from the activity tracks reported by the COVID-19 case himself/herself. Door knobs and switch buttons in room were not classified into specific room locations due to the specimens were collected with mixed form, where were commonly used by COVID-19 cases. And specimens in car (steering wheel, seat and handlebar) were also mixed form.

Air specimens was sampled using an MD8 microbiological sampler (Sartorius, Germany) and sterile gelatin filters (3 μm pores and 80 mm diameter, Sartorius, Germany)[7,14]. Air specimens was sampled for 10 minutes in toilet, anteroom and bedroom, 1 minutes in car at a speed of 50 L/minute. The filters were dissolved aseptically in 30 mL virus preservation solution (or isotonic saline solution, tissue culture solution, or phosphate buffer).

### Laboratory Procedures

The environmental surfaces and air specimens for nucleic acid detection were stored at −70 °C or below (specimens may be temporarily stored in −20 °C refrigerators in the absence of −70 °C storage condition) and were tested as soon as possible. Laboratory confirmation of environmental surfaces and air specimens was performed by qualified staff, and results were identified through open reading frame 1ab (ORF1ab) and nucleocapsid protein (N) of SARA-CoV-2 by RT-PCR testing in accordance with the protocol established by China CDC[12]. Details on laboratory processes are provided in Appendix (appendix 1 p1-2).

### Definitions

The asymptomatic persons infected with SARA-CoV-2 (asymptomatic cases in short) refers to those who have no relevant clinical manifestations including clinically detectable signs or self-perceived symptoms such as fever, cough, or sore throat, but who have tested positive for SARA-CoV-2 in respiratory specimens or other specimens[15]. Clinical classification of symptomatic cases includes 4 categories[16]: mild, moderate, severe, and critical (appendix 2 p3). In this study, the COVID-19 cases included asymptomatic, mild, moderate, severe, and critical cases. Fever was defined as an axillary temperature of 37.3°C or higher. Public area included elevator, elevator buttons, and stair armrests. An imported case is a person who was infected with SARA-CoV-2 in a foreign country and diagnosed in Guangzhou City.

### Statistical analysis

Categorical variables were presented with frequency (percentage) and continuous variables were summarized with median (interquartile range). Figures were drawn using Office Excel (version 2019). Analyses were all performed with the SAS software (version 9.4 for Windows, SAS Institute, Inc., Cary, NC, USA).

### Ethics statement

This study was based on the data from the work for an ongoing public health response to COVID-19 by GZCDC as required by the National Health Commission of China, and hence individual informed consent was waived. The study was determined not to be human subjects research and therefore was considered exempt from ethical approval after consultation with the ethics committee of GZCDC. Analytical datasets were constructed in an anonymized manner, and all analysis of personally identifiable data took place onsite at the GZCDC.

## Results

### Characteristics of 39 COVID-19 cases

Among 39 COVID-19 cases, 23 (23/39, 59.0%) were male, 11 (11/39, 28.2%) were imported and the median age was 36.0 years (interquartile range, 31.0 to 48.0 years). A total of 9 (9/39, 23.1%) asymptomatic cases and 30 (30/39, 76.9%) symptomatic cases with 10 (10/30, 33.3%) mild, 19 (19/30,63.4%) moderate, and 1 (1/30, 3.3%) severe cases were identified, and all (39/39, 100.0%) were cured and discharged. Among them, the common onset of symptoms was fever (24/39, 61.5%), dry cough (21/39, 53.9%), expectoration (9/39, 23.1%), fatigue (6/39, 15.4%), myalgia (3/39, 7.7%), diarrhea (2/39, 5.1%), and 31 (31/37, 79.5%) with CT lung abnormalities (Table 1).

**Table 1.** Characteristics of 39 COVID-19 cases in Guangzhou, China.

| Characteristics | COVID-19 cases (n = 39) |
| --- | --- |
| **Median age (IQR)** | 36.0 (31.0-48.0) |
| **Age group (%)** |  |
| <18 | 1/39 (2.5) |
| 18-44 | 22/39 (56.4) |
| 45-59 | 12/39 (30.8) |
| ≥60 | 4/39 (10.3) |
| **Male Sex (%)** | 23/39 (59.0) |
| **Imported (%)** | 11/39 (28.2) |
| **Severity (%) <sup>a</sup>** |  |
| Asymptomatic | 9/39 (23.1) |
| Symptomatic | 30/39 (76.9) |
| Mild | 10/30 (33.3) |
| Moderate | 19/30 (63.4) |
| Severe | 1/30 (3.3) |
| **Hospital admission (%) <sup>b</sup>** |  |
| Death | 0/39 (0.0) |
| Discharge | 39/39 (100.0) |
| **Highest temperature (%) (°C)** |  |
| <37.3 | 15/39 (38.5) |
| 37.3-38.0 | 19/39 (48.7) |
| 38.1-39.0 | 4/39 (10.2) |
| >39.0 | 1/39 (2.6) |
| **Symptoms at onset (%)** |  |
| Fever | 24/39 (61.5) |
| Dry cough | 21/39 (53.9) |
| Expectoration | 9/39 (23.1) |
| Fatigue | 6/39 (15.4) |
| Myalgia | 3/39 (7.7) |
| Diarrhea | 2/39 (5.1) |
| **CT lung abnormalities (%)** | **31/37 (79.5)** |
| **Median blood biochemical index (IQR)** |  |
| WBC (10<sup>9</sup>/L) <sup>c</sup> | 6.2 (4.7-7.4) |
| Ly (10<sup>9</sup>/L) <sup>d</sup> | 1.2 (0.9-1.8) |
| Ne (10<sup>9</sup>/L) <sup>e</sup> | 4.3 (3.1-5.2) |
| Ly% <sup>d</sup> | 22.6 (16.1-25.8) |
| Ne% <sup>f</sup> | 69.4 (65.7-72.5) |

a: Severity of COVID-19 at the time of diagnosis. WBC=white blood cell count; Ly=lymphocyte; Ly%=lymphocyte percentage; Ne= neutrophilic granulocyte; Ne%= neutrophilic granulocyte percentage; b: data on May 10;c: Data at hospital admission; Number of participants with missing Values: c=6, d=8, e=14, f=11.

### Distribution of environmental specimens among 39 COVID-19 cases

A total of 641 environmental surfaces and air specimens were collected among 39 COVID-19 cases, and 20 specimens (20/641, 3.1%) were positive by RT-PCR testing from 9 COVID-19 cases (9/39, 23.1%), with 5 (5/101, 5.0%) positive specimens from 3 asymptomatic cases, 5 (5/220, 2.3%) from 3 mild cases, and 10 (10/374, 2.7%) from 3 moderate cases (Table 2). The COVID-19 cases without fever (8/16 [50.0%] vs. 1/23 [4.3%]), dry cough (7/18 [38.9%] vs 2/21 [9.5%]), expectoration (8/30 [26.6%] vs. 1/9 [11.1%]), fatigue (9/33 [27.2%] vs. 0/6 [0.0%]), myalgia (20/36 [55.6%] vs. 0/3 [0.0%]), diarrhea (9/37 [24.3%] vs. 0/2 [0.0%]) symptoms were more likely to have positive specimens of SARS-CoV-2 (Table 2).

**Table 2.**
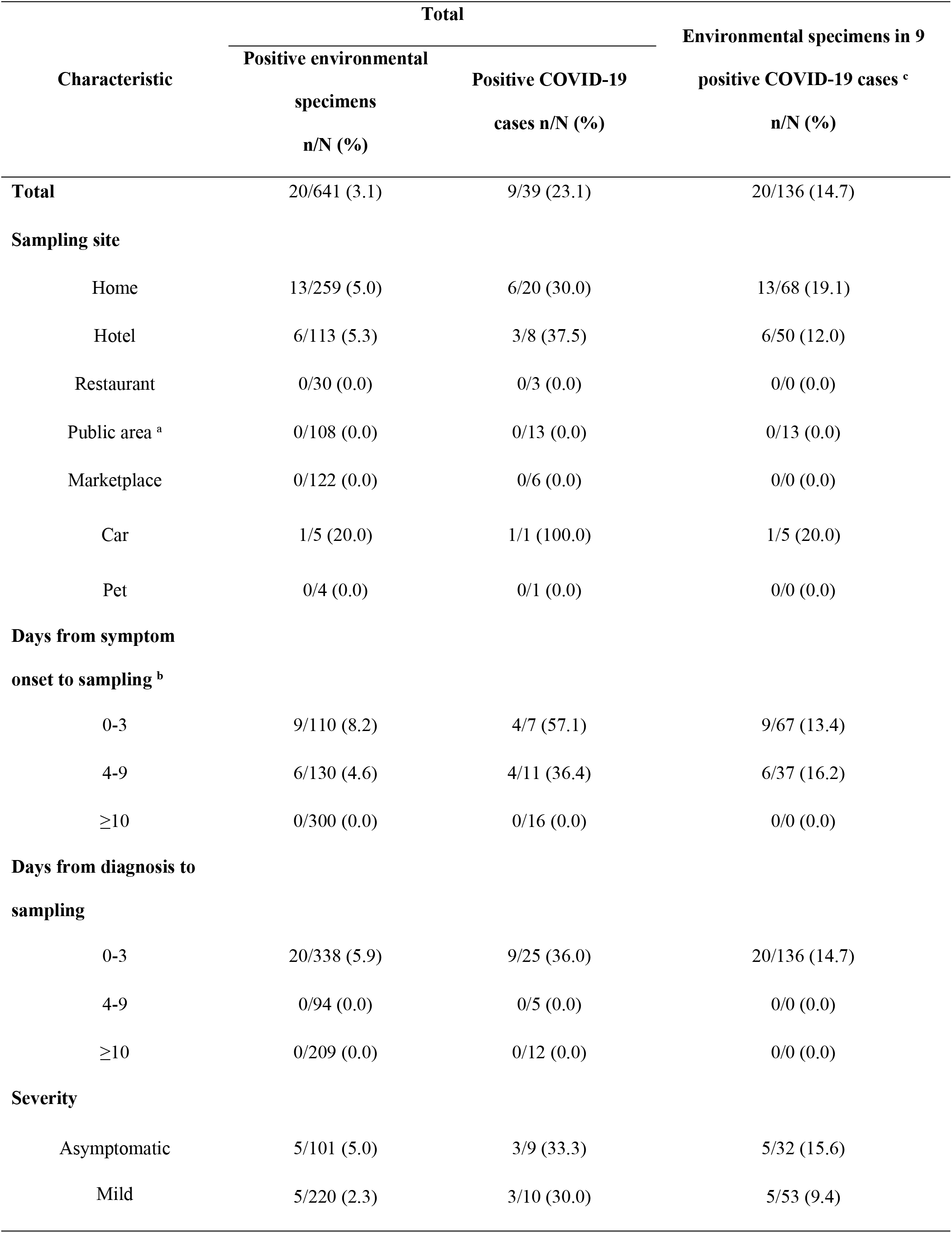
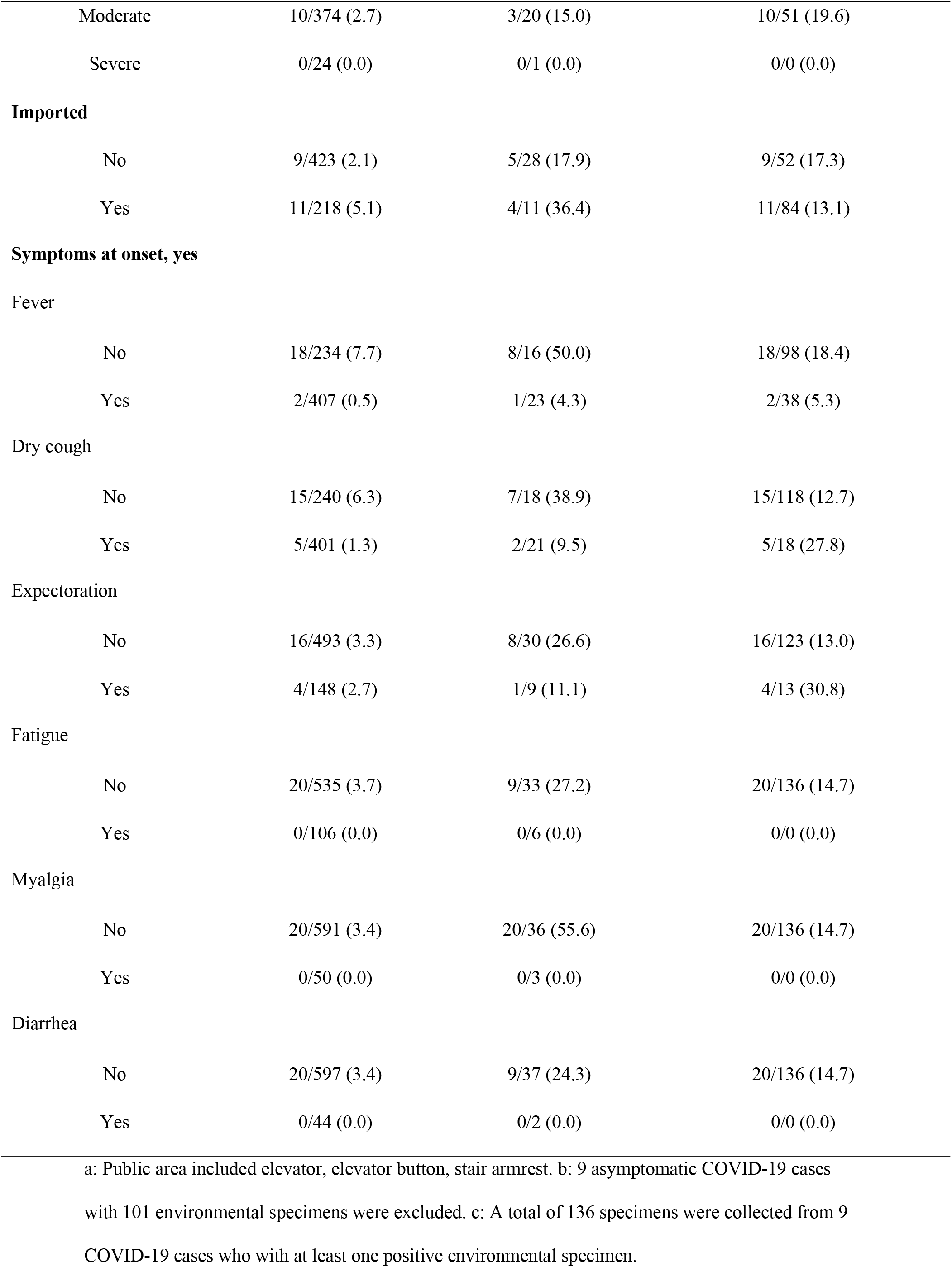
Distribution of environmental specimens among COVID-19 cases.

All the 20 positive specimens were collected within 3 days (≤3 days) from diagnosis to sampling (Figure 1), and 13 (13/259, 5.0%) positive environmental surfaces specimens were collected from home, 6 (6/113, 5.3%) from hotel and 1 (1/5, 20.0%) from car that had driven. While, specimens in restaurant that had eaten (0/30, 0.0%), marketplace that had visited (0/122, 0.0%), pet that had lived with (0/4, 0.0%) and public area that had stayed (0/108, 0.0%) were all negative by RT-PCR testing (Table 2)

**Figure 1.**
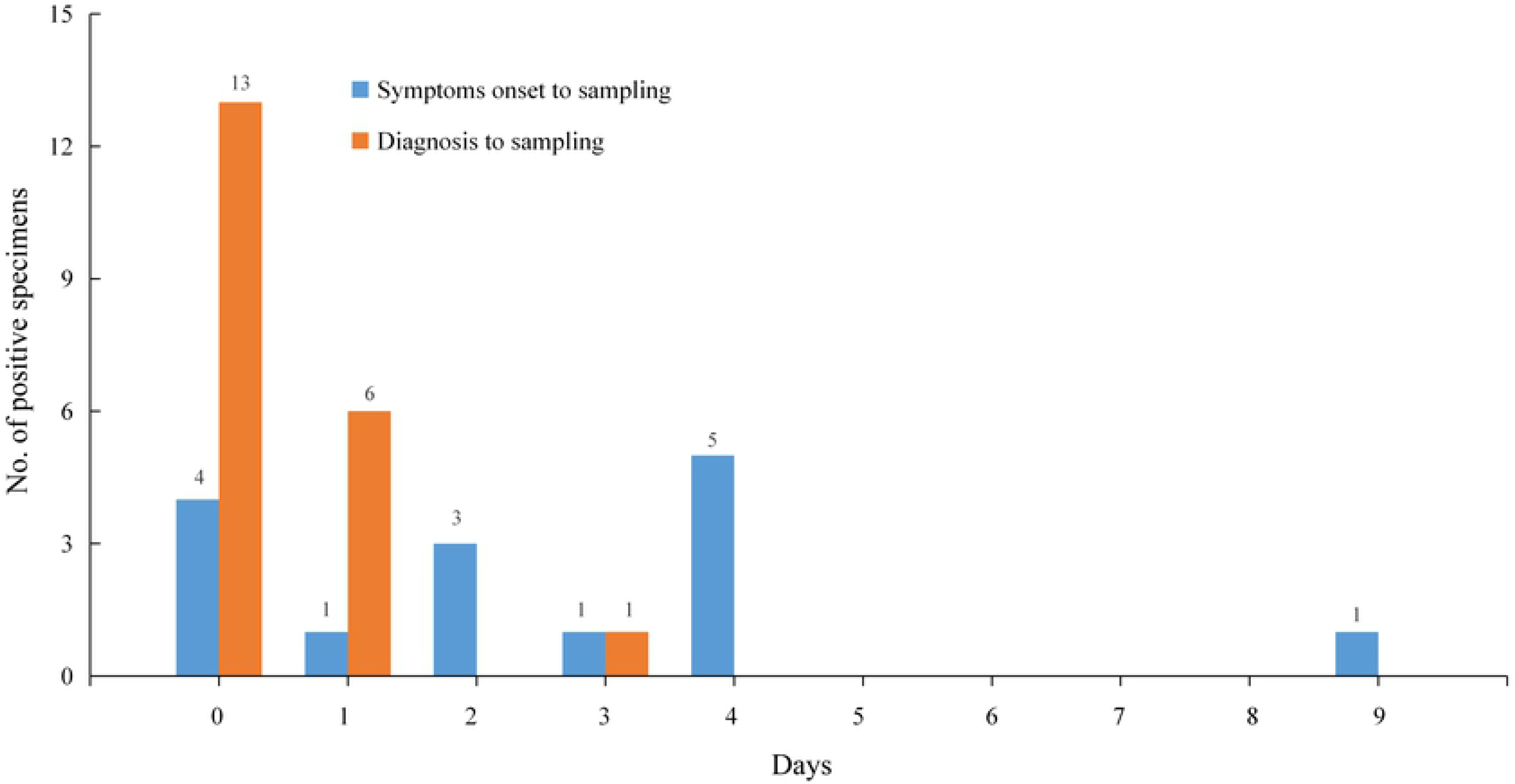
Distribution of positive environmental specimens by days from date of symptoms onset or diagnosis to sampling. (day 0 means sampling at the same day with symptom onset or diagnosis). Note: Environmental specimens were tested by RT-PCR testing; 5 positive environmental specimens of asymptomatic COVID-19 cases were excluded in bars of symptoms onset to sampling.

### Distribution of 20 positive environmental specimens among 9 COVID-19 cases

A total of 136 environmental surfaces and air specimens were collected from 9 COVID-19 cases who with at least one positive environmental specimen. Among them, 20 specimens (20/136, 14.7%) of specimens were tested positive by RT-PCR testing. Among 20 positive environmental specimens, 10 (10/42, 23.8%) were found in toilet (5 on toilet bowl, 4 on sink/faucet/shower, 1 on floor drain), 4 (4/21, 19.0%) in anteroom (2 on water dispenser/cup/bottle, 1 on chair/table, 1 on TV remote), 1 (1/8, 12.5%) in kitchen (1 on dining-table), 1 (1/18, 5.6%) in bedroom (1 on bed/sheet pillow/bedside table), 1 (1/5, 20.0%) in car (1 on steering wheel/seat/handlebar) and 3 (3/20, 21.4%) on door knobs. Surfaces specimens collected from outside room (0/13, 0.0%), air specimens collected from room (0/10, 0.0%) and car (0/1, 0.0%) were negative by RT-PCR testing (Table 3, Figure 2).

**Table 3.**
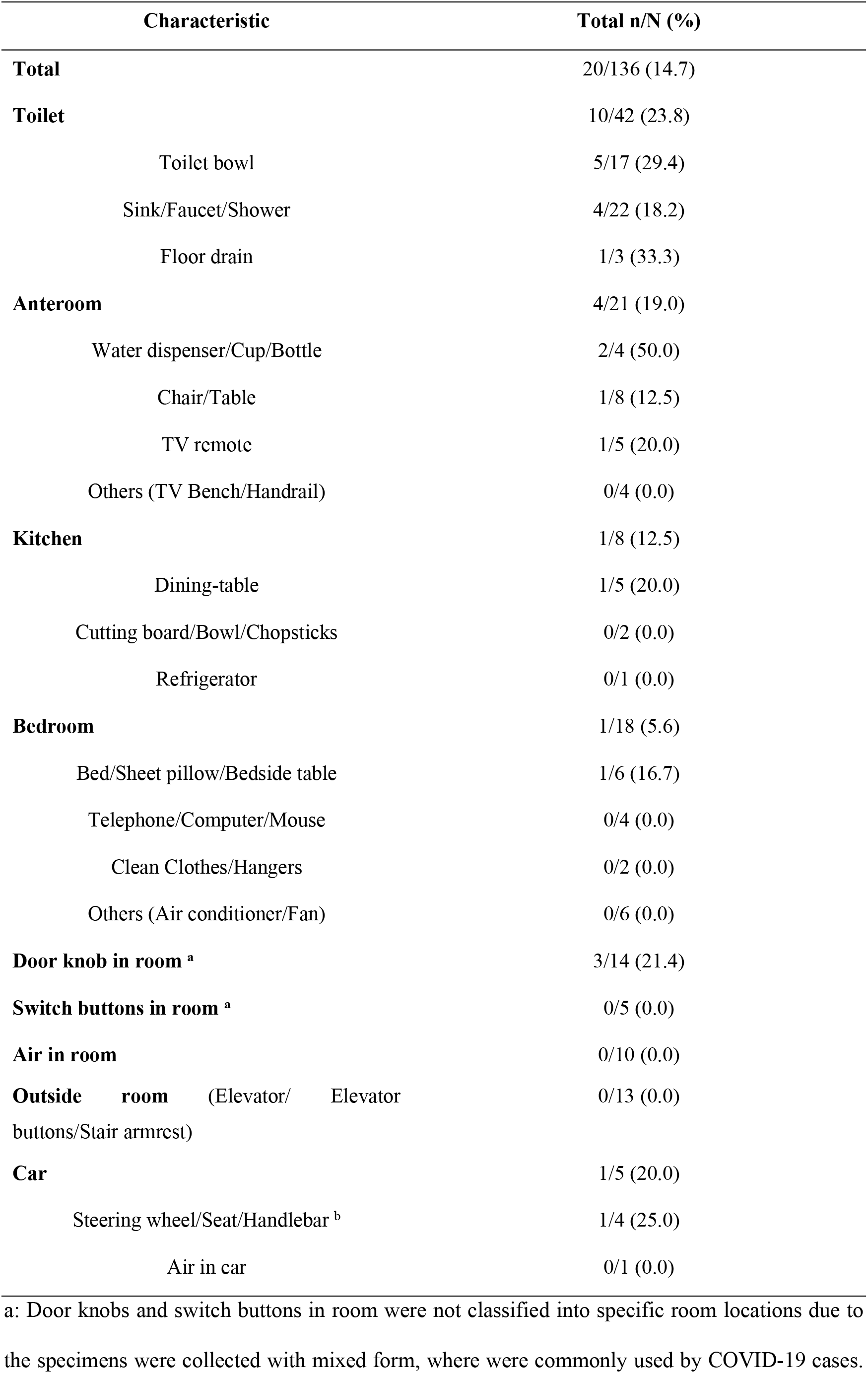
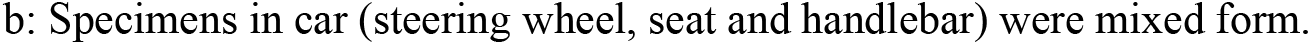
Distribution of 20 positive environmental specimens among 9 COVID-19 cases.

| **Characteristic** |  | **Total n/N (%)** |
| --- | --- | --- |
| **Total** |  | 20/136 (14.7) |
| **Toilet** |  | 10/42 (23.8) |
|  | Toilet bowl | 5/17 (29.4) |
|  | Sink/Faucet/Shower | 4/22 (18.2) |
|  | Floor drain | 1/3 (33.3) |
| **Anteroom** |  | 4/21 (19.0) |
|  | Water dispenser/Cup/Bottle | 2/4 (50.0) |
|  | Chair/Table | 1/8 (12.5) |
|  | TV remote | 1/5 (20.0) |
|  | Others (TV Bench/Handrail) | 0/4 (0.0) |
| **Kitchen** |  | 1/8 (12.5) |
|  | Dining-table | 1/5 (20.0) |
|  | Cutting board/Bowl/Chopsticks | 0/2 (0.0) |
|  | Refrigerator | 0/1 (0.0) |
| **Bedroom** |  | 1/18 (5.6) |
|  | Bed/Sheet pillow/Bedside table | 1/6 (16.7) |
|  | Telephone/Computer/Mouse | 0/4 (0.0) |
|  | Clean Clothes/Hangers | 0/2 (0.0) |
|  | Others (Air conditioner/Fan) | 0/6 (0.0) |
| **Door knob in room <sup>a</sup>** |  | 3/14 (21.4) |
| **Switch buttons in room <sup>a</sup>** |  | 0/5 (0.0) |
| **Air in room** |  | 0/10 (0.0) |
| **Outside room** (Elevator/ Elevator buttons/Stair armrest) |  | 0/13 (0.0) |
| **Car** |  | 1/5 (20.0) |
|  | Steering wheel/Seat/Handlebar <sup>b</sup> | 1/4 (25.0) |
|  | Air in car | 0/1 (0.0) |

a: Door knobs and switch buttons in room were not classified into specific room locations due to the specimens were collected with mixed form, where were commonly used by COVID-19 cases.
b: Specimens in car (steering wheel, seat and handlebar) were mixed form.

**Figure 2.**
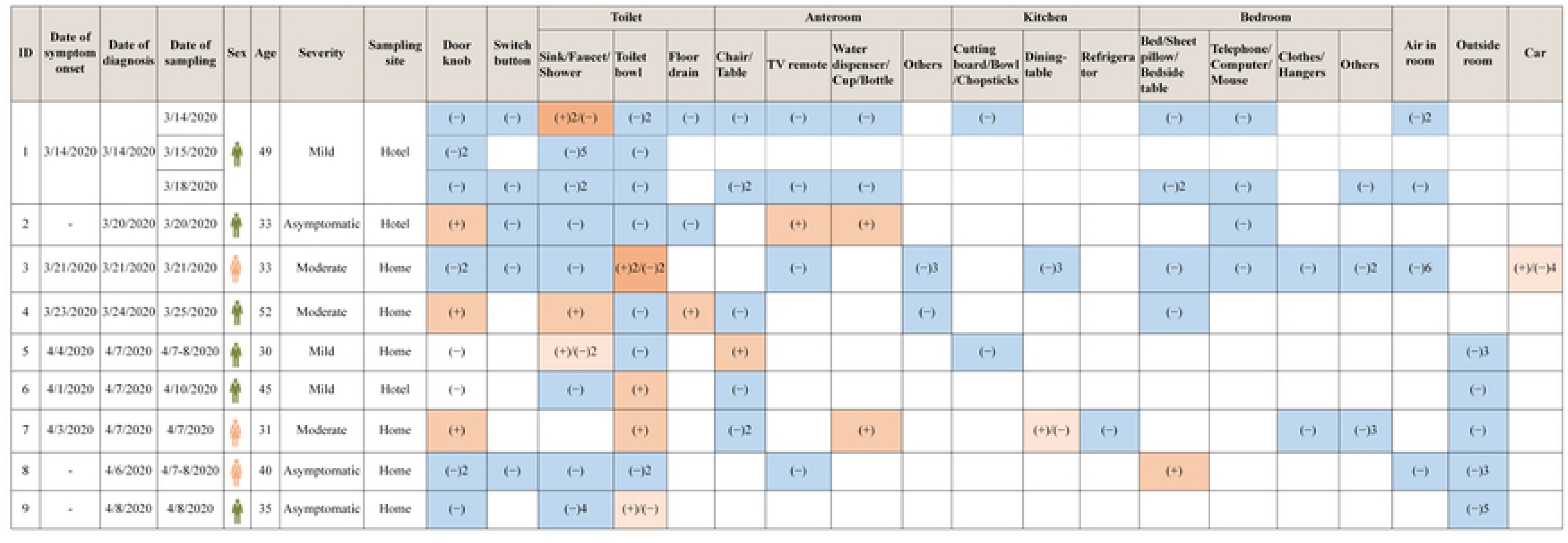
Distribution of 136 environmental specimens among 9 COVID-19 cases. Note: (+) represents positive environmental surfaces specimens, and (−) represents negative environmental surfaces and air specimens; the number represents the count of negative/positive environmental specimens; white blank represents without specimens.

## Discussion

In the current study, we evaluated the environmental contamination by SARS-CoV-2 among 641 environmental surfaces and air specimens belong to 39 COVID-19 cases before they were admitted to hospital. SARS-CoV-2 was found on environmental surfaces especially in toilet, and could survive for several days. Our study provided evidence of potential for SARS-CoV-2 transmission through contamination of environmental surfaces. All air specimens were negative. To our known, this study is the largest sample of environmental specimens with COVID-19 cases.

Previous studies showed that if one person contacted frequently with COVID-19 cases, he or she would have high incidence of infected SARS-CoV-2[3,17–19], presumably due to inhalation of droplets or contact transmission. Mathematical and animal models, and intervention researches suggested that contact transmission was the most important route in some scenarios[20]. In current study, several environmental specimens sampled from home, hotel and car were tested positive by RT-PCR testing, where were the sites that COVID-19 cases contacted frequently. Our results further demonstrated the possibility of contact transmission through environmental contamination probably due to self-inoculation of mucous membranes of mouth, nose or eyes through hands, resulting in high chance to infect SARS-CoV-2. Similar findings were made during SARS-CoV[21] and MERS-CoV[22] outbreak, in which RT-PCR positive swab specimens were confirmed from frequently touched environmental surfaces in patients’ rooms, such as medication refrigerator door, bed table and a television remote control. In other sites, such as restaurant and marketplace, there was no positive environmental specimens by RT-PCR testing in our study. However, all surrounding environmental contamination from COVID-19 cases should be alert. Although, the RT-PCR testing among specimens in restaurant and marketplace was negative, the huge number of people exposed to there also existed great risk. Previous study showed that the COVID-19 outbreak was associated with environmental contamination in restaurant[23]. In addition, some studies reported that all identified COVID-19 outbreaks of cluster confirmed cases occurred in an indoor environment[24], suggesting sharing indoor space was a major risk of SARS-CoV-2 infection.

All environmental specimens were collected before they were diagnosed in our study, which indicated that a person who exposed to environmental contamination from COVID-19 cases had a high infected risk unknowingly. In addition, SARS-CoV-2 had a great opportunity of surviving for a while on surfaces such as toilet, anteroom, and kitchen in current study. Therefore, we suggested that home quarantine for suspected COVID-19 cases might be not a good control strategy. It was difficult to ensure that cluster infection did not occur in families during the quarantine period for at least fourteen days because they shared areas like toilets during quarantine. Previous study also suggested that home quarantine required personal protective equipment and professional training, but for ordinary people and families, especially those living together in a narrow space, was obviously hard to implement excellent infection control, causing other families to be infected[11,25,26] and centralized quarantine was recommended in this condition[27].

Previous study showed that COVID-19 cases with severe disease had significantly higher viral loads than that with mild disease in respiratory specimens[28]. We tried to explore whether the more serious the COVID-19, the more contaminated to the environment surfaces. In this study, 24 environmental surfaces specimens were collected in marketplace from one severe COVID-19 cases, and the specimens’ RT-PCR testing was negative, which was probably due to the sampling site was where he or she came into contact occasionally. In other cases, several environmental surfaces specimens were tested positive with 5 (5/32, 15.6%) for 3 asymptomatic cases, 5 (5/53, 9.4%) for 3 mild cases, 10 (10/51, 19.6%) for 3 moderate cases, suggesting all cases would contaminate environmental surfaces. In addition, COVID-19 cases without symptoms like fever, dry cough, expectoration, fatigue, myalgia, diarrhea, were more likely to have positive specimens of SARS-CoV-2. It might due to that people with symptoms were quarantined more quickly in general, while, people without symptoms would continue to contaminated the environmental surfaces and air before they were admitted to hospital. We highly recommend that persons no matter COVID-19 cases or general public should conduct hand hygiene and personal protective equipment to minimize self-contamination and to protect against inoculation of mucosal surfaces and the respiratory tract[29].

Evidence suggested that influenza virus, MERS-CoV, and SARS-CoV could survive on environmental surfaces for extended periods, sometimes up to months[20,30–32]. It suggested that prolonged potential transmission of coronavirus might be existed via contact or fomite. Among environmental surfaces specimens in our study, all positive specimens were collected within 3 days (≤3 days) after diagnosis. It means that without disinfection, the SARS-CoV-2 could survive on environmental surfaces for at least 3 days, and in other studies, the survival time was 0 to 14 days[33–36], which was consistent with current understanding. Therefore, enhance surfaces cleaning and disinfection were important and essential[20].

Whether SARS-CoV-2 can be transmitted by aerosols remains controversial[5,37,38]. Fortunately, all air specimens from COVID-19 cases were negative in our study, and other studies also showed that air specimens were negative despite the extent of environmental contamination in hospital[7,37]. However, aerosol specimens in two Wuhan hospitals were tested positive collected from toilet areas, areas prone to crowding, medical staff areas[5], and suggested that SARS-CoV-2 might have potential to be transmitted via aerosols. Therefore, routine disinfection and cloth masks were recommended and some study suggested that cloth masks could potentially provide significant protection against the transmission of aerosol particles when socializing[29].

## Limitations

This study has several limitations. Firstly, the results of RT-PCR testing do not indicate the amount of viable virus, and viral culture was not done to demonstrate viability. Secondly, due to operational limitations during the outbreak, days of sampling interval was inconsistent. However, it also provided additional information about SARS-CoV-2 survival. Third, the number of air specimens represented only a small fraction of total specimens, and air exchanges in room would have diluted the presence of SARS-CoV-2 in the air. Further studies are required to confirm these preliminary results.

## Conclusions

SARS-CoV-2 was found on environmental surfaces especially in toilet, and could survive for several days. Our study provided evidence of potential for SARS-CoV-2 transmission through contamination of environmental surfaces.

## Acknowledgements

We acknowledge all staffs involved in the prevention and control of COVID-19 in Guangzhou Centers for Disease Control and Prevention.

## Data Availability

Readers interested in obtaining survey data for this study are available from CM at.

## Contribution

C.M. and D.L. contributed to the statistical analyses and had primary responsibility for writing the manuscript. L.L. and Z.Q.C directed the study. X.R.Z. and Q.M.H. contributed to the data cleaning. Z.H.L., W.Q.S. and X.F.Y. contributed to the analysis or interpretation of the data. H.Z., R.N.Z., H.P.X., J.L., J.W.L. conceived the study and supervised the collection of data. All authors critically reviewed the manuscript for important intellectual content.

